# Antibacterial Activity and Acute Toxicity Testing of Biosynthesized Silver Nanoparticles against Methicilin-Resistant *Staphylococcus aureus* and *Pseudomonas aeruginosa*

**DOI:** 10.1101/2021.06.02.446742

**Authors:** Ifeyomi Wilfred Olobayotan, Bukola Catherine Akin-Osanaiye, Olukemi A. Onuh

## Abstract

Antibacterial activity of biosynthesized silver nanoparticles was studied using the macrobroth dilution technique. The silver nanoparticles was significantly active (p > 0.05) against the test organisms at an extract concentration of 75 µg/ml. Concentrations ≤ 50 µg/ml were not as effective as the colony forming units at this concentration, 1.61 × 10^6^ for methicillin-resistant *Staphylococcus aureus* and concentrations ≤ 25 µg/ml 1.45 × 10^6^ for *Pseudomonas aeruginosa* respectively, were about the same range as the colony forming units of the controls. The silver nanoparticles inhibited Methicillin-Resistant *S. aureus* more (MIC of 75 µg/ml and MBC of 100 µg/ml) than they inhibited *P. aeruginosa* (both MIC and MBC was 100 µg/ml). The LD_50_ of the synthesized silver nanoparticles after oral administration was seen to be greater than 5000 mg/kg body weight and is therefore thought to be safe. This study supports the use of silver nanoparticles as therapeutic agents.

## INTRODUCTION

Nanoparticles are nowadays becoming very popular in various fields of research, and are useful in combating various diseases helping in early and fast detection. These particles in their pure or in the form of mixtures help in the formation of sensors, in batteries, in diagnostic kits, in water treatment, and are also helpful in curing deadly diseases such as cancer. Nanostructure materials show unique physical, chemical, biological and environmental properties, including catalytic activity, optical, electronic and magnetic properties, which have increased their applications in research, engineering, agriculture and medicine [1].

Biological silver nanoparticles (AgNPs) have the potential for large-scale applications in the dental biomaterials [2], shampoos and toothpastes [3], water purification [4] and air filtration [5]. They are also useful in clothing and textiles, medical devices and implants [6], cosmetics [7], foodstuffs packaging [8] and as effective antimicrobial agents [9]. Besides their antimicrobial properties, AgNPs have other interesting characteristics which will further enable them to be used in biosensors, electronic devices, conductive inks, catalysts and solar cells [1].

Various types of inorganic and organic nanoparticles have been utilized as antibacterial agents. The inherent antibacterial properties of some metals and metal oxides have been known for centuries. An important advantage of antibacterial metal and metal oxide nanoparticles is that they have multiple modes of action, which is why microbes can scarcely develop resistance to them. Among the inorganic antibacterial particles, silver nanoparticles are the most intensively investigated ones and capable to kill both Gram-positive and Gram-negative bacteria, having even shown to be effective against drug-resistant species [10].

Furthermore, most current antibacterial agents are chemically modified natural compounds, for instance, β-lactams (like penicillins), cephalosporins or carbapenems while some are pure natural products, such as aminoglycosides, are often used [11]. In general, these antibacterial agents can be classified as either bactericidal, which kill bacteria, or bacteriostatic, slowing down bacterial growth. Antibacterial agents are paramount to fight infectious diseases. However, with their broad use and abuse, the emergence of bacterial resistance to antibacterial drugs has become a troubling phenomenon, which is a major problem worldwide. Silver nanoparticles represent improved antibacterial agents that destroy bacteria either directly or as coupling agents that transfer drugs to target sites, without being toxic to the surrounding tissue [12]. It is therefore of importance to test the susceptibility of stubborn pathogens like multi-drug resistant *Pseudomonas aeruginosa*, a chief cause of serious nosochomial infections and methicilin-resistant *Staphylococcus aureus* responsible for a wide spectrum of infections to silver nanoparticles.

## MATERIALS AND METHODS

### Test Organisms and Standard Silver Nanoparticles

The test organisms were type cultures obtained from the diagnostic division of the National Veterinary Research Institute (NVRI), VOM, Plateau state, Nigeria. The clinical isolates of the same test organisms (for comparison) were obtainedfrom the diagnostic laboratory of University of Abuja Teaching Hospital, Gwagwalada, Abuja, Nigeria, as preserved cells on nutrient agar slants. Standard siver nanoparticles (100 µg/ml) biosynthesised using kolanut (*Cola nitida*) pod extract was obtained from LAUTECH Nanotechnology Research Group, Ladoke Akintola University, Ogbomoso, Oyo state, Nigeria.

### Media Preparation and Sterilization

All the media used were prepared according to manufacturer’s specification. Mueller-Hinton agar was used for the antibacterial testing; nutrient agar was used for the resuscitation and repeated sub-culturing of the test organisms; Mueller-Hinton broth was used for dilution broth technique; manitol salt agar was used to confirm methicillin-resistant *Staphylococcus aureus*.

### Confirmation and Purification of Test Organisms

The test organisms were resuscitated and purified by repeated sub-culturing and microscopy following gram staining.

#### Pseudomonas aeruginosa

A type culture of *Pseudomonas aeruginosa* ATCC 9027 was resuscitated on nutrient agar. Colonies produced diffusible bluish-green pigment – pyocyanin. The colonies were further gram stained and tested for chloroform solubility (by introducing an inoculum from the discrete colony in a test tube containing 10 ml of chloroform), oxidase and urease tests were also carried out for confirmation. The same procedure was repeated for the clinical isolate of the same test organism. The test organism was preserved by inoculating on nutrient agar slants in bijoux bottles incubated at 37 °C for 18-24 h and stored at 4°C until required for further use [13].

### Methicilin-Resistant *Staphylococcus aureus* (MRSA)

A type culture of methicilin-resistant *Staphylococcus aureus* ATCC6538 was first cultured on nutrient agar before it was cultured on mannitol salt agar. The growth produced yellowish colonies. This was due to the ability of *S. aureus* to use mannitol as food source leading to the production of acidic byproducts of fermentation that lower the pH of the medium and cause the pH indicator, phenol red to turn yellow. The identity and morphology of the colonies were further confirmed by microscopy following gram staining after which the cells were tested for susceptibility to methicillin. The test organism was preserved by inoculating on nutrient agar slants in bijoux bottles incubated at 37 °C for 18-24 h and stored at 4°C until required for further use. The same procedure was repeated for the clinical isolate [14].

### Standard Curve Preparation

This was done to determine the effectiveness of the silver nanoparticles synthesized as an antimicrobial agent. Standard curve were prepared using the plot of broth turbidity with a known viable colony count. The graph was then used to determine the number of viable bacteria in the broth culture with different concentrations of silver nanoparticles grown under the same conditions that the graph was created with [15]. Discrete colony of a pre-cultured methicilin-resistant *Staphyloccocus aureus* ATCC 6538 were obtained using a sterile inoculation loop and suspended in 10 ml of sterile water and the optical density was adjusted to 0.5 McFarland (≈1 × 10^8^ CFU/mL) after which the suspension was centrifuged at 2832 g for 10 minutes. The cells were washed by resuspending the sediment in clean sterile water and centrifuged at 2832 g for 5 minutes. A suspension of the cells was made by adding 1ml of the cells to 9ml of sterile peptone water and was serially diluted to obtain 10^−1^, 10^−2^, 10^−3^, 10^−4^, 10^−5^ and 10^−6^. The optical densities (OD) of the dilutions were measured using an equilibrated spectrophotometer (JENWAY 6305 UV/VIS Spectrophotometer) at 540 nm. One ml of the suspension was aseptically withdrawn from each dilution serially diluted and 0.02 ml of a chosen dilution was plated by spread plate technique on nutrient agar plates. The plates were incubated at 37 °C for 18 - 24 hours to obtain the number of colony forming units in each dilution. The calibration curve for the bacterial culture was prepared by plotting the number of cfu/ml of the dilutions against their optical densities (OD) at 540 nm. The dilution with 1 × 10^5^ CFU/ml was used as inoculum for the rest of the study (equivalent to 0.5 McFarland diluted to the ratio of 1:20 using normal saline). The same procedure was repeated for *Pseudomonas aeruginosa* ATCC9027 [16].

### Antibacterial Assay

This was carried out according to the Clinical Laboratory Standards Institute [17] methods. Broth macro-dilution method was used to assay the antibacterial activity of the AgNPs against the test organisms. Following the resuscitation on nutrient agar, a discrete colony of an 18 h culture of each type culture of the test strain was grown overnight in sterile Mueller-Hinton (MH) (HiMedia) broth on a rotary shaker (283g) at 37 °C for 18 – 24 hours. The optical density was measured and the number of colony forming units (cfu) that was inoculated deduced from the standard curve earlier prepared. A sterile Pasteur pipette was used to obtain 0.2 ml of a suspension of the test organism and was inoculated into each of the Six (6) test tubes containing 10 ml of sterile MH broth along with 0.2 ml AgNPs concentrations (5, 10, 25, 50, 75, and 100 μg/ml) respectively. The test tubes were incubated at 37 °C for 18 −24 hours. Clinically prepared antibiotics, chloramphenicol (250 mg/ml) and methicillin (250 mg/ml) were similarly tested against the test organisms under same conditions as positive controls and a test tube containing test organism with MH broth (HiMedia) as negative control. After incubation at 37 °C for 18-24 hours, the turbidity of the test tubes was visually observed and the optical densities were measured and the corresponding cfu derived from the standard curve. The assays were performed in duplicates. The same procedure was repeated for the standard silver nanoparticles obtained from the Nanotech Research Group at Ladoke Akintola University of Technology, Oyo State, tested against same test organisms under same conditions.

### Minimum Inhibitory Concentration (MIC)

The method of Vollekova [18] modified by Usman [19] for the determination of MIC was employed. In this method, the broth dilution technique where the AgNPs, the test organism and Mueller-hinton broth were added into a test tube and incubated at 37 °C for 18-24 hours. The AgNPs was prepared to the highest concentration of 100 μg/ml (stock concentration) and serially diluted (two-fold) to a working concentration ranging from 100 μg/ml down to 1.25 μg/ml. Ten ml of Mueller-hinton broth was placed in each of the different sterile test tubes and inoculated with 0.2 ml suspension of each test organism with the working concentrations of 100, 75, 50, 25, 10, 5, 2.5 and 1.25 μg/ml added to each test tube. The test tubes were incubated for 18-24 hours at 37 °C after which turbidity was observed and optical density recorded using “Cary Series UV/Vis” (Agilent Technologies, Germany). The lowest concentration that completely inhibited the growth of the test organisms as observed by the no growth when the concentration with the least optical density is plated out on nutrient agar was reported as the MIC. The assays were performed in duplicates.

### Minimum Bactericidal Concentration (MBC)

This was determined using the broth dilution resulting from the MIC tubes by sub-culturing the broth on sterile Mueller-Hinton agar as described by Vollekova [18] and Usman [19]. In this technique, inocula from those test tubes with the working concentrations where no turbidity was observed from MIC were cultured on sterile nutrient agar plates by streaking and incubated at 37 °C for 18-24 hours. The lowest concentration of the extract which showed no growth of the test organism was noted as the MBC.

### Determination of Acute Toxicity and LD_50_

The acute toxicity study was conducted according to the method of Lorke [20]. Silver is available in colloidal form. Recommended safe dose of colloidal silver for humans is 75μg/day. In the present study, different doses of silver nanoparticles were used at the time of administration for the determination of LD_50_.

### Experimental Animals

Twenty-two albino mice (35 - 60 g), male and female aged between 7-8 weeks were obtained from the Animal house of the VetMed Court Limited, Apo, Garki New Town, Abuja, Nigeria. The animals were maintained in groups of four in a metal cage at room temperature of 25 ± 2 °C and kept in the animal house in 12 hour dark/12 hour light cycle for 7 days to acclimatize. They had continuous free access to food and water during this period [21]. Each animal was used only once. For ethical reasons, all animals were sacrificed at the end of the study [22].

### Toxicological Studies in Mice

The study was conducted in two phases according to Lorke [20] using a total of twenty two male and female albino mice. In the first phase, 12 mice were divided into 3 groups of 4 mice each. Groups 1, 2 and 3 animals were given 10, 100 and 1000 mg/kg body weight (b.w.) of the silver nanoparticles, respectively, to possibly establish the range of doses producing any toxic effect. Each mouse was given a single dose after7 days of adaptation. In addition, a fourth group of four mice were set up as control group and animals in the group were not given the silver nanoparticles.

In the second phase, further specific doses (1500, 3000 and 5000 mg/kg b.w.) of the silver nanoparticles were administered to six mice (two mice per dose) to further determine the LD_50_ value. The silver nanoparticles was dissolved in Phosphate buffered saline (PBS) solution and given via oral route. All animals were observed frequently (after 2, 4, 6 and 8 hours) on the day of treatment for any change in behaviour or physical activities and surviving animals were monitored daily for 2 weeks for signs of acute toxicity. Recovery and weight gain were seen as indications of having survived the acute toxicity. At the end of 14 days, all surviving mice were sacrificed with Pentothal sodium (40 mg/kg) and then autopsied.

### Calculation of Median Lethal Dose (LD_50_)

For each mouse, the observation was made for 14 days and symptoms of toxicity and rate of mortality in each group were noted. At the end of study period, expired animals were counted for the calculation of LD_50_. The arithmetic method of Lorke [20] was used for the determination of LD_50_.

LD_50_= √ (D_0_ x D_100_)

D_0_ = Highest dose that gave no mortality,

D_100_ = Lowest dose that produced mortality.

### Statistical Analysis

The statistical analyses were carried out using statistical package for social sciences (SPSS). A simple t – test of hypothesis was conducted to determine whether there was variation in the antibacterial activity of the silver nanopartcles to the significance level of 5%. Values of the effect of silver nanoparticles on the body weight of the mice during acute toxicity testing were compared using the analysis of variance (ANOVA). For all analyses the level of statistical significance was fixed at p≤0.05.

## RESULTS

The antibacterial testing was carried out in duplicates for each concentration, hence, the colony forming units were recorded per concentration with the average and standard deviation recorded (Table 1). Chloramphenicol was not tested against methicillin-resistant *S. aureus*, and methicillin was not tested against *P. aeruginosa*. From the table, concentrations ≥75 µg/ml showed pronounced inhibitory activity against the test organisms.

**Table 1:**
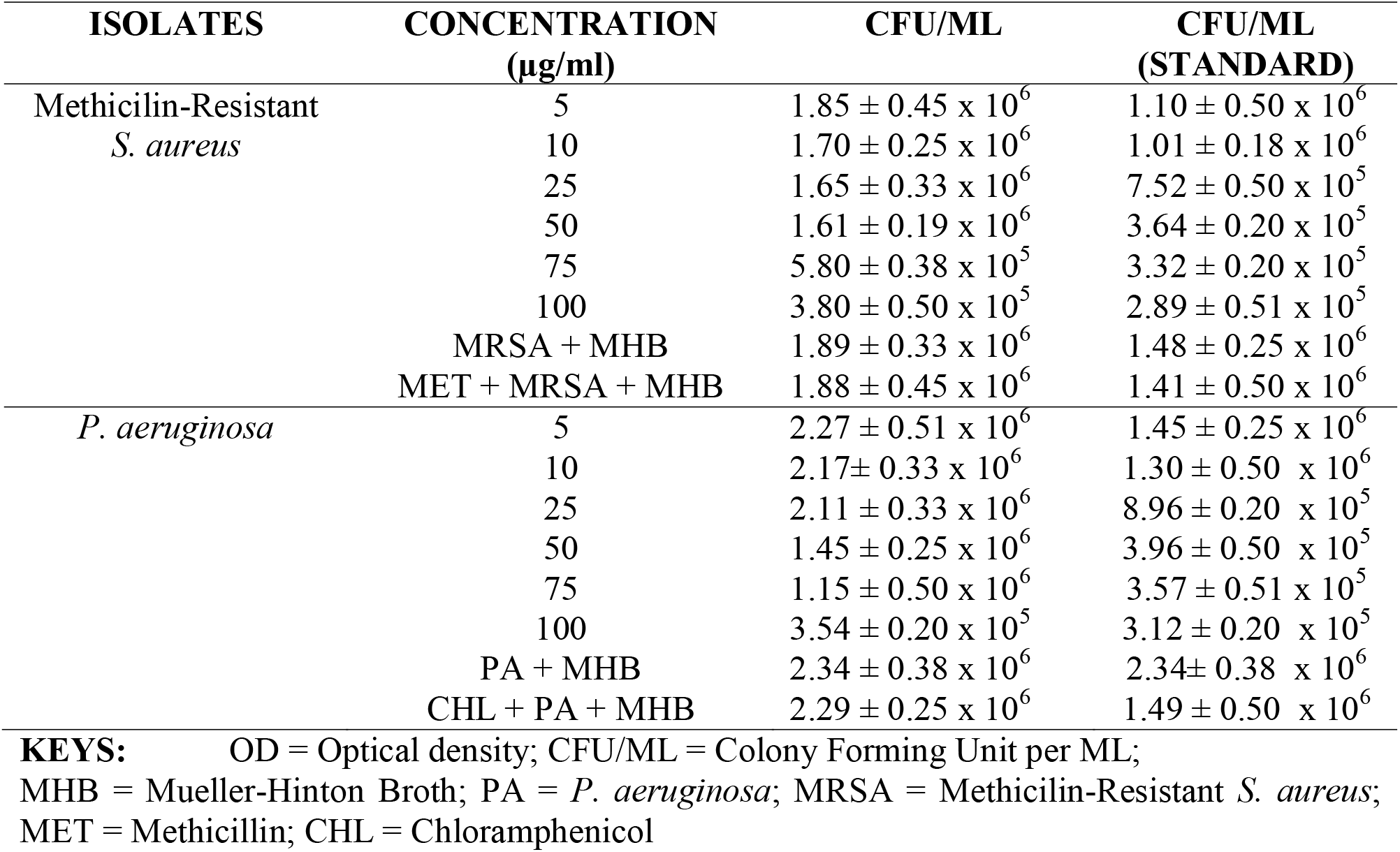
Antibacterial Activity of Synthesized AgNPs by Macrobroth Dilution.

| ISOLATES | CONCENTRATION<br>($\mu\text{g/ml}$) | CFU/ML | CFU/ML<br>(STANDARD) |
| --- | --- | --- | --- |
| Methicilin-Resistant<br>*S. aureus* | 5 | $1.85 \pm 0.45 \times 10^6$ | $1.10 \pm 0.50 \times 10^6$ |
| | 10 | $1.70 \pm 0.25 \times 10^6$ | $1.01 \pm 0.18 \times 10^6$ |
| | 25 | $1.65 \pm 0.33 \times 10^6$ | $7.52 \pm 0.50 \times 10^5$ |
| | 50 | $1.61 \pm 0.19 \times 10^6$ | $3.64 \pm 0.20 \times 10^5$ |
| | 75 | $5.80 \pm 0.38 \times 10^5$ | $3.32 \pm 0.20 \times 10^5$ |
| | 100 | $3.80 \pm 0.50 \times 10^5$ | $2.89 \pm 0.51 \times 10^5$ |
| | MRSA + MHB | $1.89 \pm 0.33 \times 10^6$ | $1.48 \pm 0.25 \times 10^6$ |
| | MET + MRSA + MHB | $1.88 \pm 0.45 \times 10^6$ | $1.41 \pm 0.50 \times 10^6$ |
| *P. aeruginosa* | 5 | $2.27 \pm 0.51 \times 10^6$ | $1.45 \pm 0.25 \times 10^6$ |
| | 10 | $2.17 \pm 0.33 \times 10^6$ | $1.30 \pm 0.50 \times 10^6$ |
| | 25 | $2.11 \pm 0.33 \times 10^6$ | $8.96 \pm 0.20 \times 10^5$ |
| | 50 | $1.45 \pm 0.25 \times 10^6$ | $3.96 \pm 0.50 \times 10^5$ |
| | 75 | $1.15 \pm 0.50 \times 10^6$ | $3.57 \pm 0.51 \times 10^5$ |
| | 100 | $3.54 \pm 0.20 \times 10^5$ | $3.12 \pm 0.20 \times 10^5$ |
| | PA + MHB | $2.34 \pm 0.38 \times 10^6$ | $2.34 \pm 0.38 \times 10^6$ |
| | CHL + PA + MHB | $2.29 \pm 0.25 \times 10^6$ | $1.49 \pm 0.50 \times 10^6$ |

**KEYS:** OD = Optical density; CFU/ML = Colony Forming Unit per ML; MHB = Mueller-Hinton Broth; PA = *P. aeruginosa*; MRSA = Methicilin-Resistant *S. aureus*; MET = Methicillin; CHL = Chloramphenicol

Table 2 shows the least concentration of the silver nanoparticles that had inhibitory effect on the test isolates. From the table, the silver nanoparticles inhibited Methicillin-Resistant *S. aureus* more (MIC of 75 µg/ml) than they inhibited *P. aeruginosa* (MIC of 100 µg/ml).

**Table 2:**
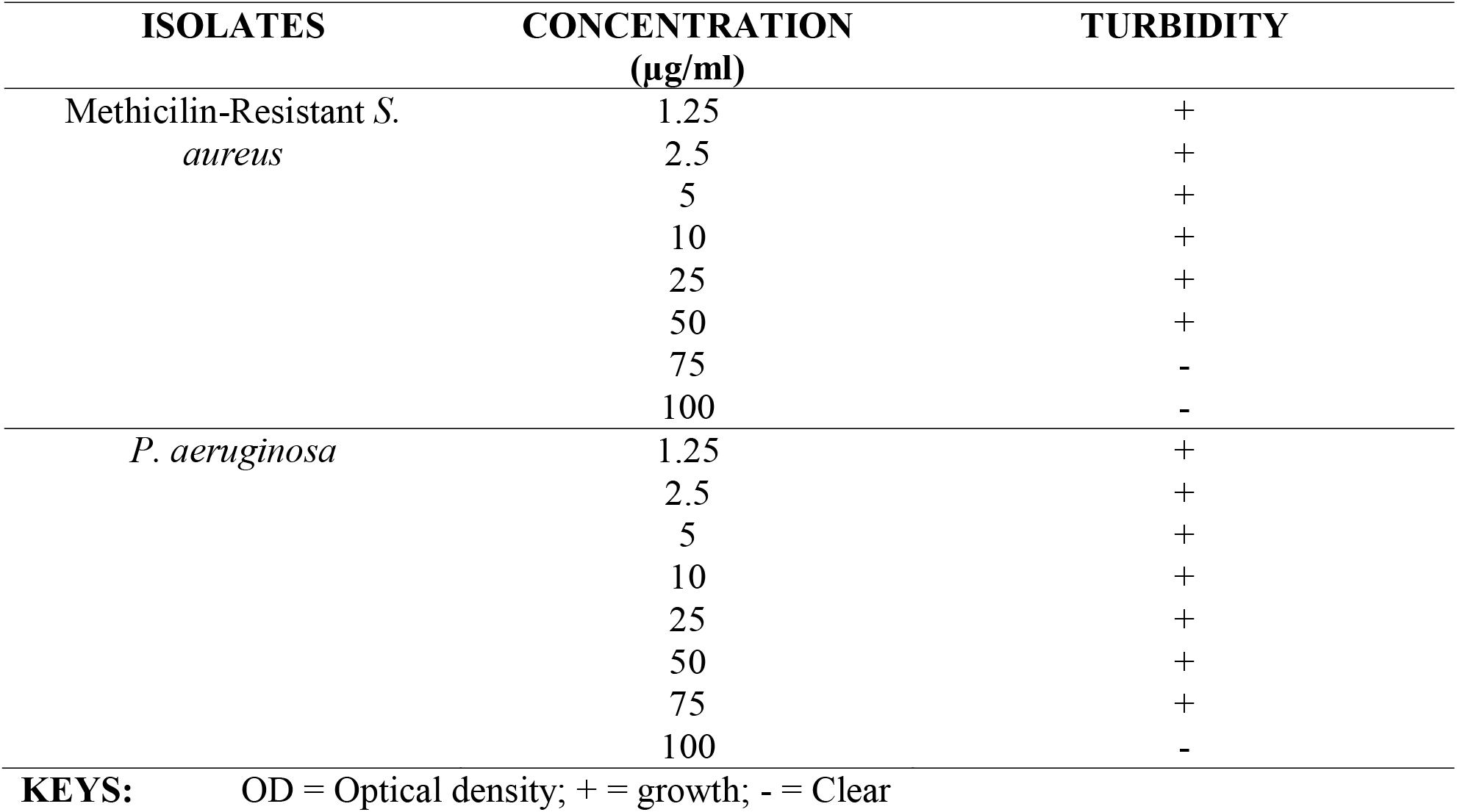
Minimum Inhibitory Concentration of the Synthesized AgNPs by Macrobroth Dilution.

Table 3 shows the least concentration of the synthesized silver nanoparticles that had a lethal effect on the test isolates. It was seen that at a concentration ≤ 75 µg/ml, the test isolates could still survive but at a concentration ≥ 100 µg/ml, the test isolates could not withstand the effect of the silver nanoparticles. The minimum bacterial concentration (MBC) is 100 µg/ml for both test isolates.

**Table 3.**
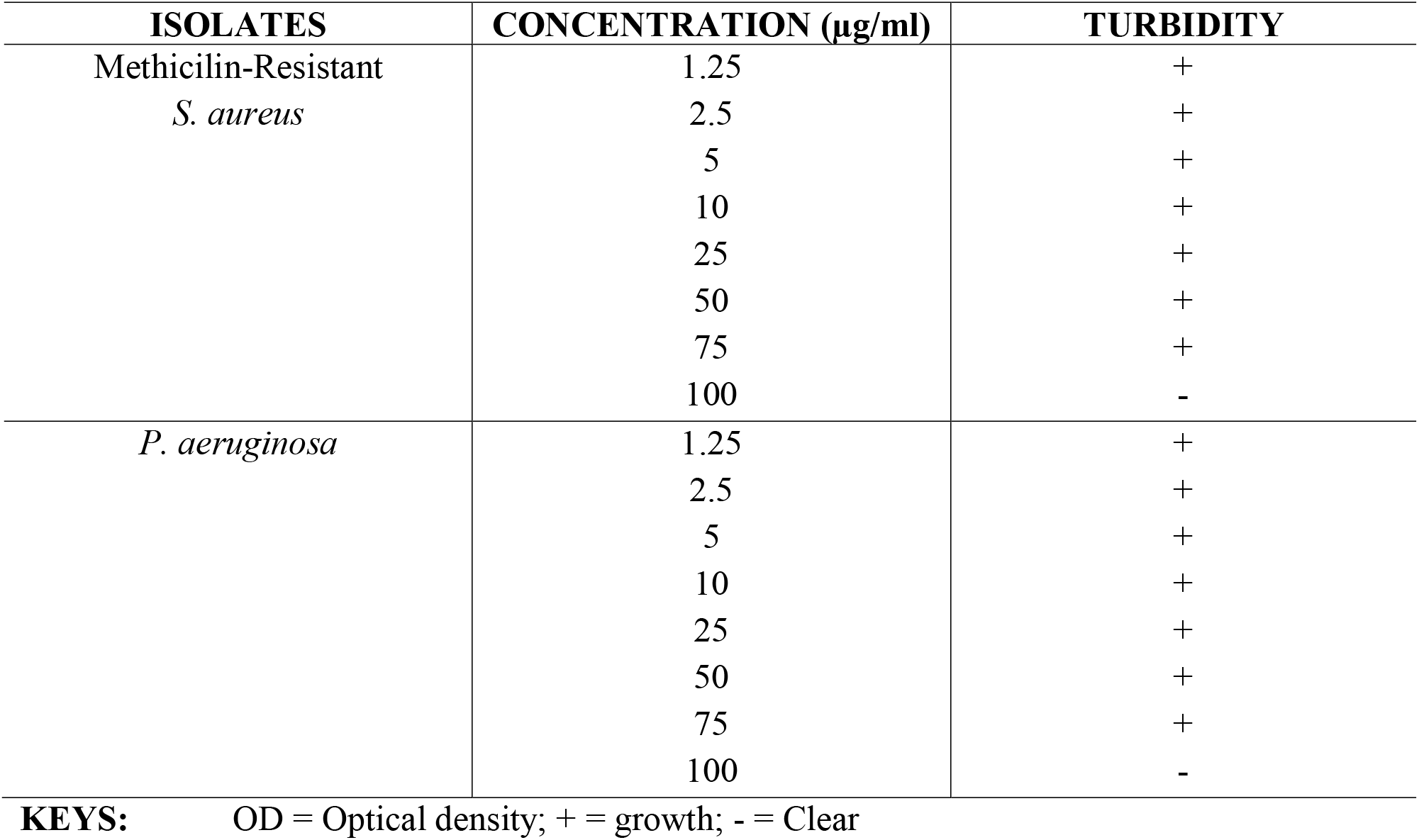
Minimum Bactericidal Concentration of the Synthesized AgNPs by Macrobroth Dilution.

| ISOLATES | CONCENTRATION ($\mu\text{g/ml}$) | TURBIDITY |
| --- | --- | --- |
| Methicilin-Resistant<br>*S. aureus* | 1.25 | + |
|  | 2.5 | + |
|  | 5 | + |
|  | 10 | + |
|  | 25 | + |
|  | 50 | + |
|  | 75 | + |
|  | 100 | - |
| *P. aeruginosa* | 1.25 | + |
|  | 2.5 | + |
|  | 5 | + |
|  | 10 | + |
|  | 25 | + |
|  | 50 | + |
|  | 75 | + |
|  | 100 | - |

Table 4 shows the acute lethal effect of the silver nanoparticles on albino mice.

**Table 4.** Acute Toxicity Testing of Silver Nanoparticles on Albino Mice.

| **Experiment** | **Dose (mg/kg bw)** | **No. of dead rats**<br>**after 24 hours** | **No. of dead rats**<br>**after 14 days** |
| --- | --- | --- | --- |
| Phase 1 | 10 | 0/3 | 0/3 |
|  | 100 | 0/3 | 0/3 |
|  | 1000 | 0/3 | 0/3 |
| Control | 0 | 0/3 | 0/3 |
| Phase 2 | 1500 | 0/3 | 0/3 |
|  | 3000 | 0/3 | 0/3 |
|  | 5000 | 0/3 | 0/3 |

From the table, it can be seen that no albino mice died after 24 hours of oral administration of the silver nanoparticles and there was also no casualty after the 14 days duration of the experiment. Therefore, it is observed that the LD_50_is >5000 mg/kg.

Table 5 shows the effect of oral administration of the silver nanoparticles on the body weight of rats during acute toxicity.

**Table 5:**
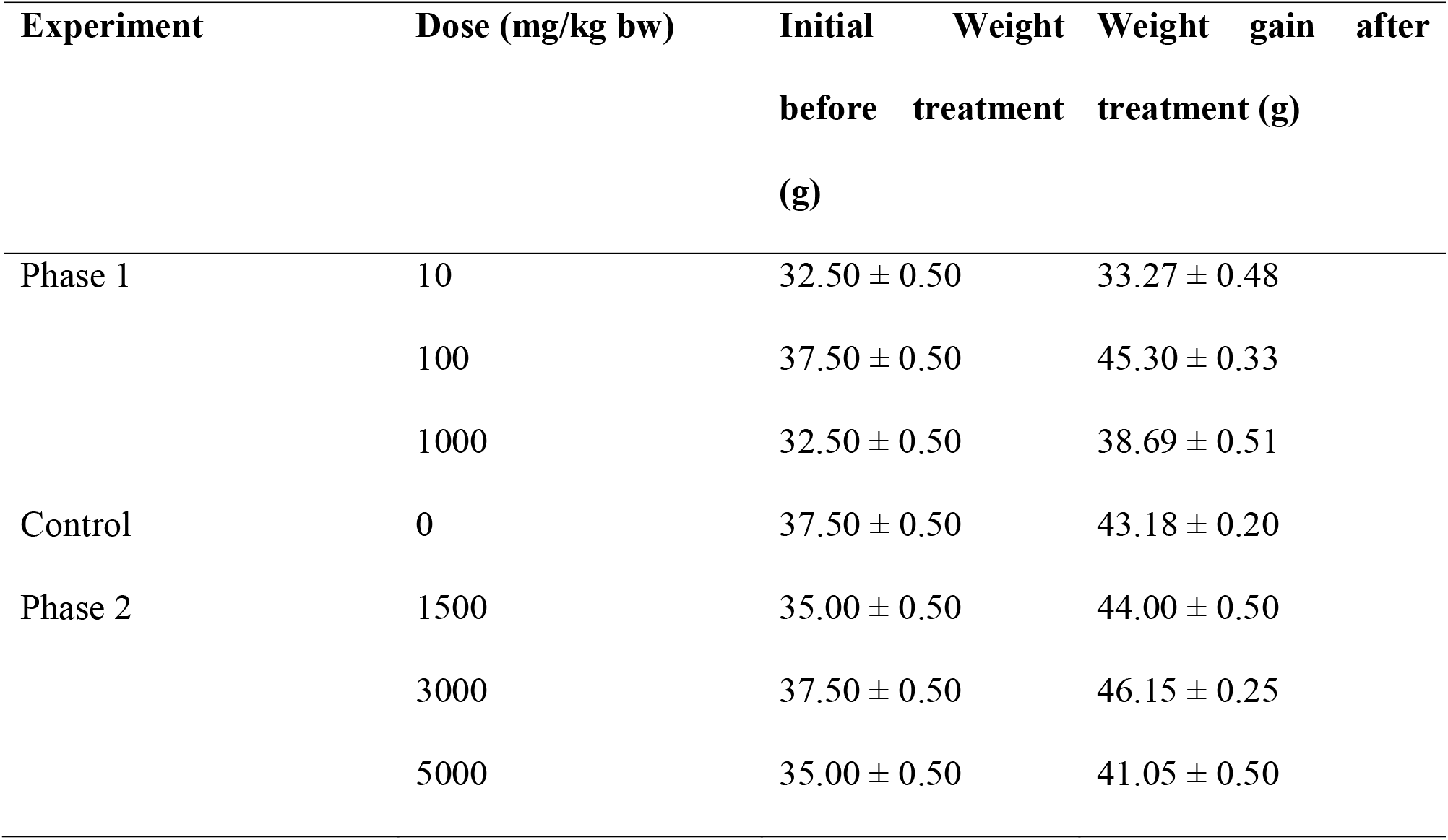
Effect of Oral Administration of Silver Nanoparticles on the body weight of albino mice.

From the table above, it can be seen that the mice gained weight after oral administration of the silver nanoparticles which was not found to be significant (p>0.05) when compared with the control group.

Table 6 shows the some pathology result for acute toxicity of silver nanoparticles orally administered to albino mice after they were sacrificed.

**Table 6:** Post Mortem Result for Acute Toxicity of Silver Nanoparticles Orally Administered to Albino Mice.

| Gross Pathology Observe |  |  |  |  |
| --- | --- | --- | --- | --- |
| Organ | Dose (mg/kg bw) |  |  |  |
| Liver | Normal | Normal | Normal | Normal |
| Lungs | Normal | Normal | Normal | Normal |
| Kidneys | Normal | Normal | Normal | Normal |
| Heart | Normal | Normal | Normal | Normal |
| Spleen | Normal | Normal | Normal | Normal |

From the table, it was observed macroscopically that some of the internal organs of the mice were normal, therefore the silver nanoparticles is thought to be non-toxic at the dose administered.

## DISCUSSION

The antibacterial susceptibility pattern of the silver nanoparticles (Table 1 – 3) showed the silver nanoparticles was significantly active (p > 0.05) against the test organisms at an extract concentration of 75 µg/ml. Concentrations ≤50 µg/ml were not as effective as the colony forming units at this concentration, 1.61 × 10^6^ for methicillin-resistant *Staphylococcus aureus* and concentrations ≤25 µg/ml 1.45 × 10^6^ for *Pseudomonas aeruginosa* respectively were about the same range as the colony forming units of the controls. Chloramphenicol was not tested against methicillin-resistant *S. aureus*, and methicillin was not tested against *P. aeruginosa*. The activity of the synthesized silver nanoparticles against both Gram -positive and Gram-negative bacteria is an indication of its broad spectrum antibacterial activity. Results obtained showed that the silver nanoparticles were more active against *S. aureus* the Gram-positive organism than *P. aeruginosa* the Gram-negative organism. This could be ascribed to the morphological differences between these microorganisms. Gram-negative bacteria have an outer phospholipids membrane carrying the structural lipopolysaccharide components, which makes the cell wall partially impermeable. Gram-positive bacteria however, have only an outer peptidoglycan layer which is not an effective permeability barrier and are thus more susceptible [23].

The activity of the synthesized silver nanoparticles was found to be less effective than the standard nanoparticles obtained from the nanotech research laboratory in Lautech (Table 1), which showed profound activity against both test organism at a concentration of ≥10 µg/ml (1.01 × 10^6^ and 1.30 × 10^6^ colony forming units for methicillin-resistant *Staphylococcus aureus* and *Pseudomonas aeruginosa* respectively) and conforms with the findings of Wen-ru *et al*. [24] who worked on the antibacterial activity and mechanism of silver nanoparticles on *Escherichia coli* with 10 µg/ml effectively inhibiting the growth of the test organism. The reduced activity of the synthesized silver nanoparticles may be due to the presence of impurities that may like interfere with its mode of action of inhibition of cell membrane synthesis of the test organisms.

The silver nanoparticles inhibited Methicillin-Resistant *S. aureus* more (MIC of 75 µg/ml and MBC of 100 µg/ml) than they inhibited *P. aeruginosa* (both MIC and MBC at 100 µg/ml). It was observed that the values of the MIC and MBC of the synthesized silver nanoparticles on methicillin-resistant *S. aureus* were different while the values on *P. aeruginosa* were the same (Tables 2 and 3). Hugo and Rusell [25] reported that with most bactericidal antimicrobials, the MIC and MBC are often near or equal in values. Therefore, it can be said that the silver nanoparticles had a bacteriostatic effect on methicillin-resistant *S. aureus* and a bactericidal effect on *P. aeruginosa*, which indicates that the silver nanoparticles can help in the treatment of respiratory tract infections especially in individuals with a compromised respiratory tract or a compromised systemic defense mechanism. In a similar research, Wen-ru *et al*. [24] reported that the MIC and MBC of silver nanoparticles was 10 µg/ml, when tested against *Escherichia coli* while Ayala-Nunez *et al*. [25] reported that the MIC and MBC of silver nanoparticles was 13.5 µg/ml, when it was tested against methicillin-resistant *Staphylococcus aureus*.

The results revealed that the synthesized silver nanoparticles were not found to be toxic after 24 hours of oral administration at 5000 mg/kg body weight (Table 4). The albino mice that received the 5000 mg/day body weight orally survived and non-significant changes were observed. However, there was slight change in behavioural pattern as seen by the corner sitting, salivation and general weakness noticed. The albino mice were autopsied and there were no physically observed changes in the vital organs. The LD_50_ value (Table 6) of the synthesized silver nanoparticles after oral administration being greater than 5000 mg/kg body weight is therefore thought to be safe as suggested by Lorke [20]. Furthermore, the non-mortality of the mice throughout the two weeks of the experiment also agrees with this claim.

Statistical analysis showed that the potency of the silver nanoparticles against both test organisms was not significantly different from each other (p > 0.05). The effect of the doses of the silver nanoparticles administered to the mice was not found to be statistically significant on their weight.

## CONCLUSION

In this study, it has been demonstrated that silver nanoparticles are effective antibacterial agents as shown by its inhibition of the pathogenic test organisms. It was also noted that the silver nanoparticles, when used in at required quantities are not toxic to This study provides some scientific basis of the use of silver nanoparticles in medicine as antimicrobial agents and for targeted drug delivery, in food sector as preservatives and in agriculture to treat plant diseases.

## ACKNOWLEDGEMENT

I appreciate the efforts of the staff of Nanotech Research Group, Ladoke Akintola University, Ogbomoso, Nigeria, who provided standard silver nanoparticles for comparison and the management and staff of Vetmed Hub Limited, Apo, Abuja, Nigeria for providing needed assistance in making the research work possible.

## REFERENCES

[1] Khatami, M., Pourseyedi, S., Khatami, M., Hamidi, H., Zaeifi, M., and Soltani, L. and (2015). Synthesis of Silver Nanoparticles using Seed Exudates of Sinapis arvensis as a Novel Bioresource, and Evaluation of their Antifungal Activity. Bioresources and Bioprocessing, 2: 1–7.

[2] Corrêa, J. M. (2015). Silver Nanoparticles in Dental Biomaterials. International Journal of Biomaterials, 1: 1–9.

[3] Bansod, S D., Bawaskar, M S., Gade, A K., and Rai, M. K. (2015). Development of Shampoo, Soap and Ointment formulated by Green Synthesised Silver Nanoparticles Functionalised with Antimicrobial Plants Oils in Veterinary Dermatology: Treatment and Prevention Strategies. Institute of Engineering, Technology and Nanobiotechnology, 9: 165–171.

[4] Dankovich, T. A. (2014). Microwave-assisted Incorporation of Silver Nano particles in Paper for Point-of-Use Water Purification. Environmental Science: Nano, 1: 367–378.

[5] Ko, Y., Joe, Y. H., Seo, M., Lim, K., Hwang, J., and Woo, K. (2014). Prompt and Synergistic Antibacterial Activity of Silver Nanoparticle-decorated Silica Hybrid Particles on Air Filtration. Journal of Material Chemistry B, 2: 6714–6722.

[6] De Mel, A., Chaloupka, K., Malam, Y., Darbyshire, A., Cousins, B., and Seifalian, A. M., (2012). A Silver Nanocomposite Biomaterial for Blood-contacting Implants. Journal of Biomedicine and Material Resource Part A100A, 9: 2348–2357.

[7] Kokura, S., Handa, O., Takagi, T., Ishikawa, T., Naito, Y., and Yoshikawa, T. (2010). Silver Nanoparticles as a Safe Preservative for Use in Cosmetics,” Nanomedicine: Nanotechnology, Biology and Medicine, 6: 570–574.

[8] Becaro, A A., Puti, F C., Correa, D S., Paris, E C., Marconcini, J M., and Ferreira, M. D. (2015). Polyethylene Films Containing Silver Nanoparticles for Applications in Food Packaging: Characterization of Physico-Chemical and Antimicrobial Properties. Journal of Nanoscience and Nanotechnology Research, 15: 2148–2156.

[9] Patil, S. S., Dhumal, R. S., Varghese, M. V., Paradkar, A. R. and Khanna, P. K. (2015). Nanoparticles. Synthesis and Reactivity in Inorganic, Metal-Organic and Nano-Metal Chemistry, 39: 65.

[10] Zhu, X., Radovic-Moreno, A. F., Wu, J., Langer, R., and Shi, J. (2014). Nanomedicine in the Management of Microbial Infection – Overview and Perspectives. Nano Today, 9(4):478–98.

[11] von Nussbaum, F. (2006). Antibacterial Natural Products in Medicinal Chemistry – Exodus or Revival? Angewandte Chemie International Edition, 45:5076–5079.

[12] Witte, W. (2004). International Dissemination of Antibiotic Resistant Strains of Bacterial Pathogens. Infection, Genetics and Evolution, 4:187–191.

[13] Sandeep, K. S. (2010). Isolation and Characterisation of Pseudomonas. In: Detection and Analysis of Tuberculosis. Germany: Lambert Academic Publishers. Pg. 200–204.

[14] Clauditz A., Resch A., Wieland K. P., Peschel A., Götz F. (2006). Staphyloxanthin Plays a Role in the Fitness of Staphylococcus aureus and its Ability to Cope with Oxidative Stress. Infection and immunity, 74 (8): 4950–4953.

[15] Vlab, A. (2011). Bacterial Growth Curve. Biotechnology and Biomedical Engineering, 5–7.

[16] Isu, N. R. and Onyeagba, R. A. (1999). Basic Practicals in Microbiology. 1^st^ Edition. Okigwe: Fasmen Communications. Pg. 202–203.

[17] Clinical and Laboratory Standards Institute. (2007). Performance Standards for Antimicrobial Susceptibility Testing; Seventeenth Informational Supplement (M100-S17); Wayne, PA: Clinical and Laboratory Standards Institute. Pg. 25.

[18] Vollekova, A.D., Kostalova, V. and Sochorova, R. (2001). Isoquinoline Alkaloids from Mahonia aquifolium stem bark is active against Malassezia Spp. Folia Microbiology, 46:107–111.

[19] Usman, H., Abdulrahman, F. I. and Ladan, A. H. (2007). Phytochemical and Antimicrobial Evaluation of Tribulus terrestis L. (Zygophylaceae) Growing in Nigeria. Research Journal in Biological Science Medwell Journals, 293):244–247.

[20] Lorke, D. (1983). A New Approach to Practical Acute Toxicity Testing. Archives of Toxicology, 53:275–287.

[21] Organisation for Economic Co-operation and Development (OECD), (2001). Guidelines for Testing of Chemicals. Acute Oral Toxicities up and down Procedure. 425, 1–26. www.oecd.org/dataoecd/17/51/1948378.pdf

[22] American Veterinary Medical Association (AVMA) Guidelines for the Euthanasia of Animals: 2013 Edition. AVMA, United States, Pg. 50.

[23] Nostro, A., Germano, M. P., D’Angelo, V., Marino, A. and Cannatelli, M. A. (2000). Extraction methods and bioautography for evaluation of medicinalplant antimicrobial activity. Letters of Applied Microbiology, 30: 379–384.

[24] Wen-ru, L., Xiao-Bao, X., Qing-Shan, S., Hai-Yan, Z., You-Sheng, O. U. and Yi-Ben, C. (2010). Antibacterial Activity and Mechanism of Silver Nanoparticles on Escherichia Coli. Applied Microbiology and Biotechnology, 85: 1115–1122.

[25] Hugo, W. B. and Rusell, A. D. (2011). Pharmaceutical Microbiology. In: Denyer, S. P., Hodges, N., Gorman, S. P. and Gilmore, B. (eds). 8th Edition. UK: Blackwell 7. Science Ltd. Pp 229–245.

[26] Ayala-Nunez, V. N., Humberto, H. L., Liliana, I. T. and Cristina, R. P. (2009). Silver Nanoparticles Toxicity and Bactericidal Effect Against Methicillin-Resistant Staphylococcus aureus: Nanoscale Does Matter. Nanobiotechnology, 7: 7–6.

